# Synergistic effects of combing proton- or X-irradiation with anti-PDL1 immunotherapy in two murine oral cancers

**DOI:** 10.1101/2023.09.13.557140

**Authors:** Anne Marit Rykkelid, Priyanshu Manojkumar Sinha, Charlemagne Asonganyi Folefac, Michael Robert Horsman, Brita Singers Sørensen, Tine Merete Søland, Olaf Joseph Franciscus Schreurs, Eirik Malinen, Nina Frederike J Edin

## Abstract

**Background and purpose:** Combining radiation therapy with immunotherapy may be beneficial in treatment of head and neck cancer (HNC), but the combined effect may depend on tumor characteristics and the type of radiation. The purpose was to compare responses for two syngeneic tumor models in mice following X-ray or proton irradiation with or without immune checkpoint inhibition (ICI).

**Materials and methods:** MOC1 and MOC2 tumors were inoculated in the right hind leg of each mouse (C57BL/6J, n=159). Single-dose irradiation with X-rays or protons and administration of anti-PDL1 started when the tumors reached 200 mm^3^. Doses of 5-30 Gy were given. Time-dependent tumor volume data were analyzed with a regression model yielding the growth rate γ without irradiation and the reduction in growth rate per dose η. Relative biological effectiveness (RBE) was calculated as the ratio of η for X-rays to that of protons. Synergy between radiation and ICI was estimated as the ratio of η’s.

**Results:** MOC2 tumors grew faster and were more radioresistant than MOC1 tumors. ICI reduced the growth rate for MOC1 with 20±2% compared to controls, while no reduction was seen for MOC2. RBE for MOC1 wo/w ICI was 0.89±0.04 and 0.93±0.06, respectively, while it was 1.15±0.12 and 1.60±0.17, respectively, for MOC2. Combination synergy for X-rays was 1.22±0.08 and 0.96±0.11 in MOC1 and MOC2, respectively, while was it 1.27±0.06 and 1.33 ±0.13, respectively, for protons.

**Conclusions:** RBE for protons was dependent on use of ICI and tumor type. A greater synergy may be achieved when combining protons with ICI compared to X-rays and ICI.

## Introduction

Radical radiotherapy of head and neck cancer (HNC) has largely remained unchanged during the last decades, with survival levels for patients with human papilloma virus (HPV) negative disease at around 50% [1]. Intensity modulated radiotherapy (IMRT) using high-energy X-rays has reduced the prevalence and severity of side effects compared to 3D conformal radiotherapy [2], but proton therapy may further reduce toxicities [3]. Compared to X-rays, protons have higher relative biological effectiveness (RBE), which currently is set to a constant of 1.1 in clinical practice [4]. However, RBE varies with proton energy and is likely dependent on tissue type and endpoint [4]. Moreover, it is an open question whether RBE changes when combining proton therapy with systemic therapies [5].

Programmed cell death protein 1 (PD-1) is an inhibitory receptor expressed on several types of immune cells, which upon interaction with programmed death-ligand 1 (PDL1) is activated and inhibit cytokine secretion, T cell activation, and T cell proliferation and survival. Tumor cells may escape discovery and elimination by overexpression of PDL1, thus limiting the immune response [6]. Immunotherapy has emerged as a new pillar in cancer treatment, and use of immune checkpoint inhibitors (ICIs) such as anti-PDL1 has been shown to significantly prolong the survival of cancer patients across several cancer types [7-9]. It has also shown an ability to suppress tumor growth in several different murine tumor models [10-12]. However, the majority of patients with HNC do not respond to ICI monotherapy [13], exemplified by a recent a study which found a response rate of 13 % to anti-PD1-treatment [14]. This can be due to lack of anti-tumor T cell activation and infiltration and/or the presence of immunosuppressive cells and cytokines.

There is increasing interest in combining ionizing radiation with immunotherapy to improve outcomes, as radiation may cause immunomodulatory effects [15]. Ionizing radiation may promote the release of damage associated molecular patterns (DAMPs) that in turn attract and activate anti-tumor T cells [16, 17]. However, irradiation may at the same time trigger immunosuppressive responses [18]. Protons and other ions have been suggested to be superior to X-rays in stimulating the immune system due to the differences in energy deposition patterns compared to X-rays, which has been shown for a colon cancer mouse model [19]. Still, the immunostimulatory and immunosuppressive actions of ionizing radiation may be dependent on dose, fractionation, and radiation quality [20], and the mechanisms dependencies and impact on treatment outcomes are poorly understood.

To our knowledge, the efficacy of combined proton therapy and ICI, and comparison with X-ray therapy, has not been previously investigated. Also, RBE estimates for protons in combination with ICI are generally lacking. In this study, we have assessed the effect of anti-PDL1 immunotherapy in combination with proton or X-irradiation in C57BL/6 mice with tumors from two murine oral cancer cell lines with different immunogenicity; MOC1 and MOC2. The aim was to determine if the combination of proton therapy and ICI gave a synergistic effect compared to proton therapy alone, and to compare this to X-rays with or without ICI.

## Method

### Murine mouse model

Two Mouse Oral Cancer (MOC) cell lines were obtained (Kerafast, USA). MOC1 is a cell line with indolent growth *in vivo* and it is very immunogenic, while MOC2 is a less immunogenic, aggressive cancer with rapid growth *in vivo* and propensity to metastases [21, 22]. C57BL/6Jrj female mice were used throughout (Janvier, France). The mice were 10-12 weeks (first round of experiments) or 16-18 weeks (second and third round of experiments) at inoculation. Tumor tissue was first implanted into the mouse flank. When the tumor reached a size of approximately 1000mm^3^ it was harvested and processed into a cell suspension in PBS, and 5-8 μl was injected subcutaneously into the right hind leg of each mouse. At least 7 mice were included in each treatment group (Table 1, supplementary material).All animal experiments were performed in accordance with the animal welfare policy of Aarhus University and approved by the Danish Animal Experiments Inspectorate.

**Table 1:** Growth rate (γ) and radiation-induced reduction in growth rate (η) with ±standard error and 95 % CI for protons and X-rays alone or in combination with anti-PDL1-treatment for MOC1 and MOC2 tumors. Results stem from fitting the regrowth delay model (eq. 1) to time-dependent tumor volume data. The goodness of fit is indicated by the coefficient of determination (R^2^).

| Treatment groups | Growth rate, $\gamma$ | Growth rate reduction, $\eta$ | $R^2$ |
| --- | --- | --- | --- |
| **<u>MOC1</u>** |  |  |  |
| Protons | 0.073±0.001 [0.072, 0.075] | 0.0041±0.001 [0.0038, 0.0044] | 0.69 |
| X-rays |  | 0.0046±0.002 [0.0043, 0.0050] | 0.75 |
| Protons + anti-PDL1 | 0.065±0.001 [0.058, 0.062] | 0.0052±0.001 [0.0048, 0.0056] | 0.57 |
| X-rays + anti-PDL1 |  | 0.0056±0.002 [0.0051, 0.0061] | 0.60 |
| **<u>MOC2</u>** |  |  |  |
| Protons | 0.200±0.003 [0.195, 0.205] | 0.0030±0.002 [0.0025, 0.0034] | 0.64 |
| X-rays |  | 0.0026±0.002 [0.0022, 0.0030] | 0.64 |
| Protons + anti-PDL1 | 0.202±0.004 [0.196, 0.209] | 0.0040±0.003 [0.0034, 0.0045] | 0.62 |
| X-rays + anti-PDL1 |  | 0.0025±0.002 [0.0021, 0.0029] | 0.62 |

### Treatment and follow-up

Treatment started when the tumor size reached approximately 200 mm^3^ (range 150-257 mm^3^). Tumor volume (TV) was calculated as an ellipsoid volume from length (L), width (W) and height (H) caliper measurements as TV = π/6 x L x W x H. TV was measured every weekday after treatment until end of follow up. Mice were euthanized when the tumor reached 1000mm^3^ or three times TV compared to day 0, or earlier if humane endpoint was reached. At euthanization, the tumor was harvested (when possible) and immediately put in formalin buffer for fixation with duration of 24-72H at 5 degrees Celsius before it was dehydrated and embedded in paraffin.

After initial ICI screening with anti-CTLA4, anti-PD1, and anti-PDL1, the latter was chosen as it resulted in the largest partial growth inhibition for MOC1 tumors. In the current study, mice were injected intraperitoneally with anti-PDL1 (InVivoMAb anti-mouse PDL1 (B7-H1), Clone 10F.9G2, BioXCell, USA), stock concentration 7.95 mg/ml, 10mg/kg mouse or with PBS on days 1, 4, 8 and 11 after irradiation.

Irradiations with both X-rays and protons were performed using the same setup. Non-anesthetized mice were restrained in a custom-designed Lucite jig, with the right foot fixed in an outstretched position on a resting plate attached to the jig. It was ensured that the foot was not clamped to avoid reduced blood supply and hypoxia. The mice were placed in a custom water tank with the foot fully emerged in water with temperature 25°C during irradiation. The mouse body was shielded with a brass collimator and only the tumor-bearing leg was in the radiation field. For proton irradiations, the leg was placed in the middle of the spread-out Bragg peak using a horizontal scanning proton pencil-beam (ProBeam, Varian, USA). The proton irradiation procedure has been described in detail elsewhere[23, 24]. X-irradiations were performed using a YXLON Maxishot X-ray unit (280kV, 2Gy/min) with the dosimetry controlled by a Semiflex ionization chamber (PTW, Germany). The doses given to the tumor in a single fraction at day 0 were 5, 10, 15, and 20 Gy for MOC1 and 10, 20, and 30 Gy for MOC2.

All short-term TV data were analyzed using a regression approach (see Data analyses below). Mice with MOC2 tumors were euthanized due to metastases typically 6-19 days after treatment, and it was not possible to access long-term effects. Response categories were defined in order to estimate long-term treatment effect for MOC1 tumors. Cutoff was chosen at day 45 after treatment for MOC1 tumors. The categories were: (1, CR) Complete Remission = No trace of primary tumor, (2, PR) Partial Remission = Primary tumor smaller than at start of treatment, (3, TR) Temporary Response = Primary tumor shrinks to smaller size than at start of treatment and stays below that size for 5 days or more but with significant regrowth, and (4, PD) Progressive disease = Mouse lives shorter than the cutoff day and/or has a continually growing primary tumor.

### Data analyses

All analyses were done in Python v3.8.8 with libraries NumPy, SciPy, and statsmodels. Error bars represent 1 standard error of the mean (SEM). The correlation *r* between two series was calculated as Pearson’s correlation coefficient. Significance was defined as no overlap between 95% confidence intervals. Goodness of fit was given by the coefficient of determination (R^2^-value).

To estimate both the impact of anti-PDL1 treatment and radiation dose on tumor growth, a simple mathematical model was developed. The model assumes that without irradiation, the tumor volume increases exponentially with time explained by a growth rate *γ* (in unit of [day^-1^]). Upon irradiation, the growth rate is reduced with the delivered radiation dose *D* by a proportionality constant *η* [Gy^-1^ day^-1^]. This constant reflects the tumor radiosensitivity in terms of growth rate reduction per dose. Thus, the model can be expressed as:

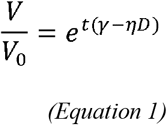

where *V* and *V*_0_ is tumor volume on day *t* and zero, respectively. The model is expected to perform well on short-term volume data where the irradiation inactivates a certain fraction of the tumor cells, which lowers the apparent growth of the bulk tumor. The model was fitted to all data in given treatment group (X-rays±anti-PDL1, protons±anti-PDL1) with non-linear regression. Moreover, as *η* implicitly reflects tumor radiosensitivity, RBE was estimated using:

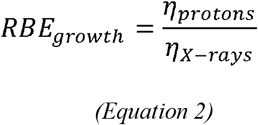

With corresponding synergy estimated for each radiation type as:

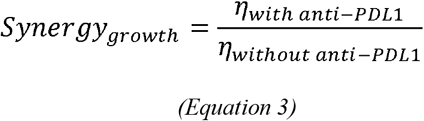

Tumor response data in terms of binary treatment effect were analyzed using logistic regression and subsequent estimate of the tumor dose giving 50 % effect (TD_50_). Effect was defined as permanent or temporary remission (CR, PR or TR) while no effect was defined as progressive disease (PD). RBE of binary treatment effect (TE) was calculated as:

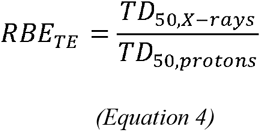

Correspondingly, the synergy was estimated for each radiation type as:

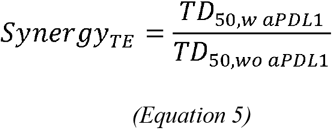

where w/wo aPDL1 is with/without anti-PDL1 administration.

## Results

MOC1 and MOC2 cells showed very different tumor growth characteristics when implanted in the mice. MOC2 tumors grew to approximately 200mm^3^ (treatment size) in about 6-19 days after inoculation with a median of 8 days, while for MOC1 it took about 15-68 days after inoculation (median of 28 days). Ninety-two % of MOC1 and all MOC2 inoculated mice successfully developed tumor in the right hind foot. For MOC1, a few instances of metastasis occurred in the lymphatic system. For MOC2, all mice developed lymph node or lung metastases and had to be euthanized between day 3 and 25 (median 12 days) even if the primary tumor was reduced by the treatment.

For MOC1 tumors, anti-PDL1 treatment alone significantly reduced tumor growth rate γ with 20±2% (fig.1a, 3a, table 1), while for MOC2 there was no effect of anti-PDL1 treatment alone (fig.2a, 3c, table 1). To access the radiation-induced reduction in growth rate, the data were fitted to the growth model (eq. 1, fig.1 and 2). The model performance was good with R^2^>0.57 and small uncertainties in estimated model parameters (table 1). The cutoffs for the analysis were chosen at day 19 (MOC1) and day 7 (MOC2) post initiation of treatment, as these were the latest time points where all treatment groups had at least 80% surviving animals.

**Figure 1:**
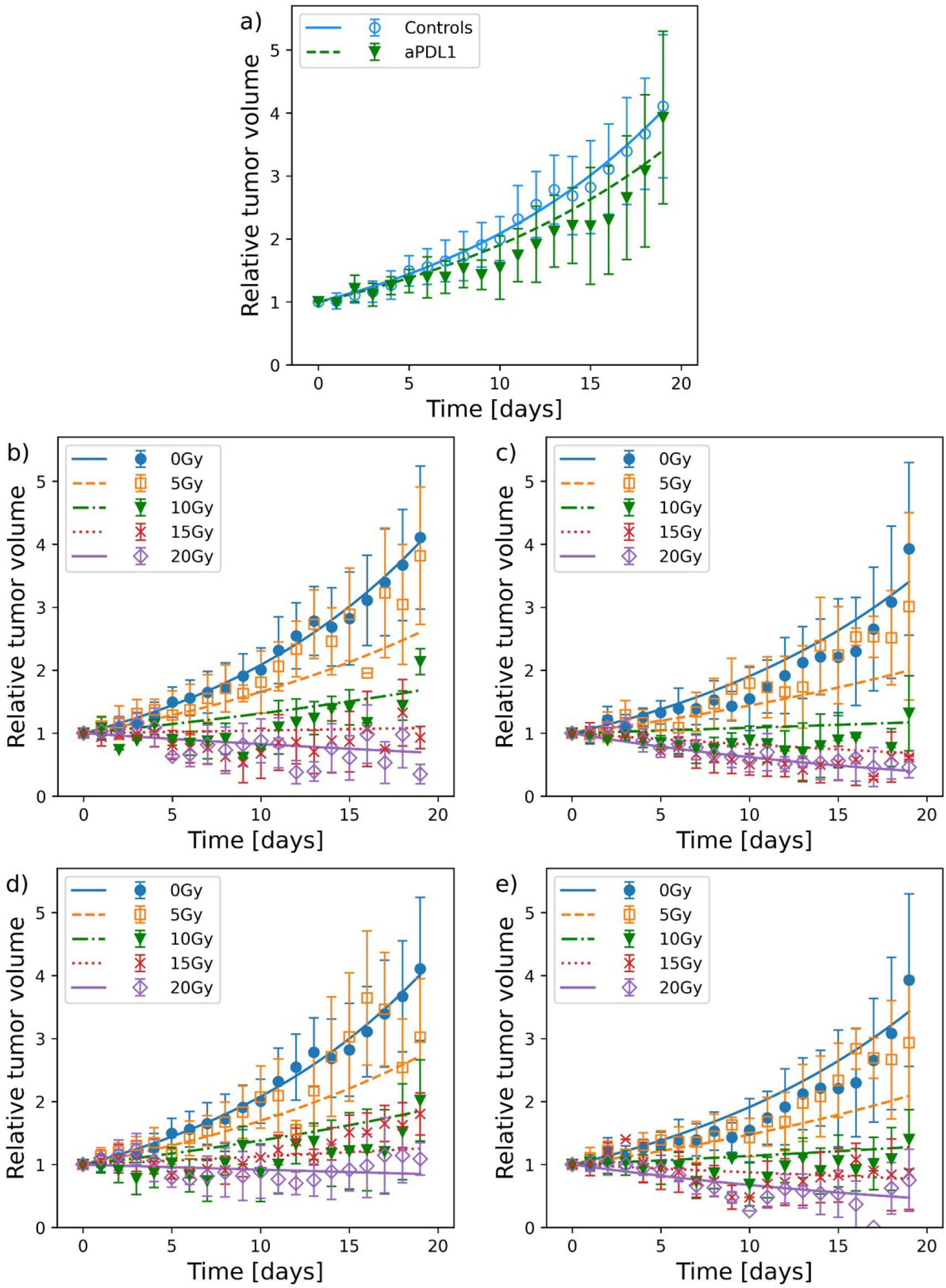
Relative volume plotted against time for MOC1 tumors (a-e) with lines corresponding to model fits (eq. 1) with resulting growth rate γ and growth rate reduction per dose η (Fig.3a-b). a) Unirradiated tumors with or without anti-PDL1, b) X-irradiation, c) X-irradiation + anti-PDL1, d) Proton irradiation, e) Proton irradiation + anti-PDL1. Error bars correspond to 1 SEM.

**Figure 2:**
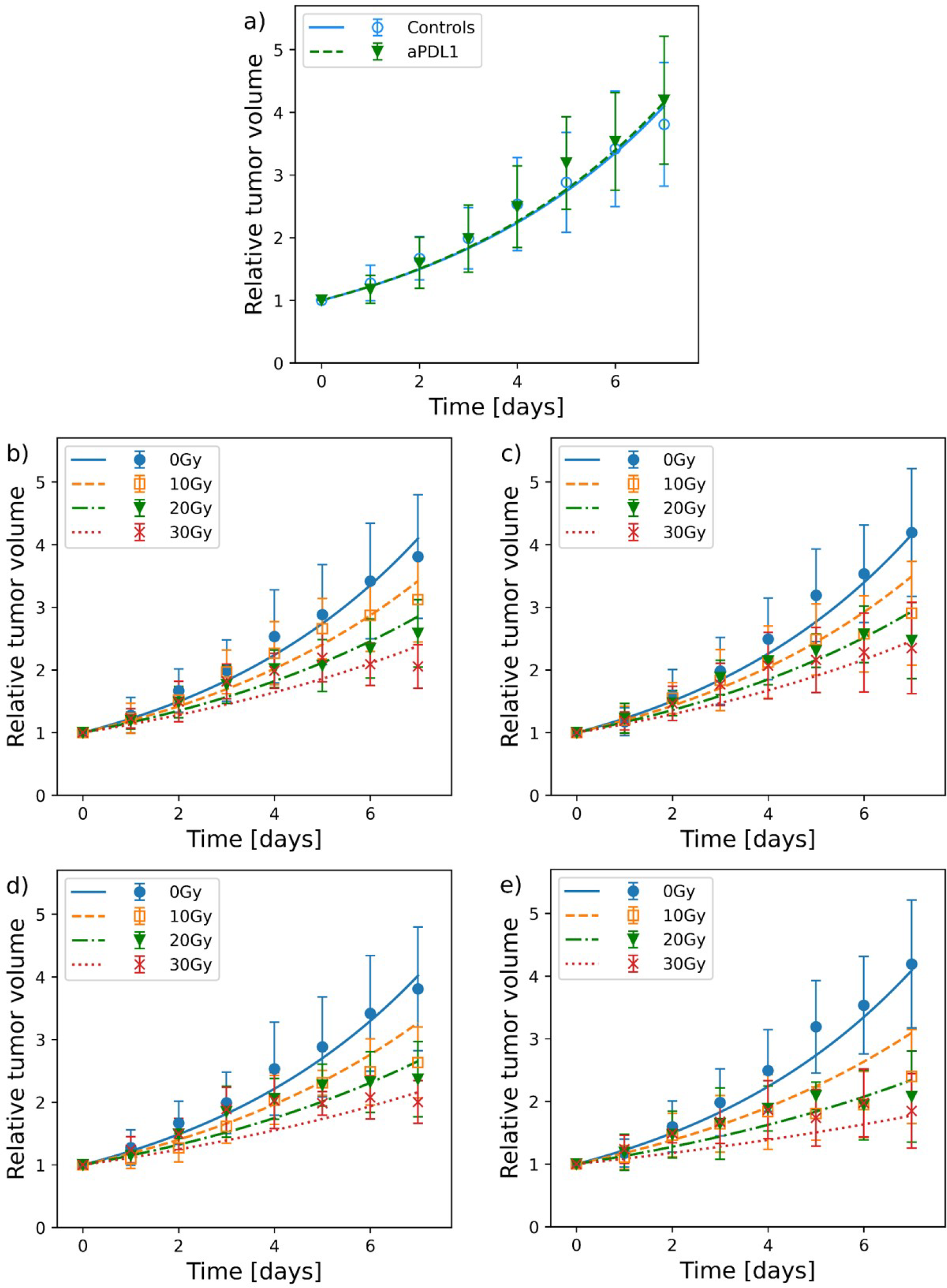
Relative volume plotted against time for MOC2 tumors (a-e) with lines corresponding to model fits (eq. 1) with resulting growth rate γ and growth rate reduction per dose η (Fig.3c-d). a) Unirradiated tumors with or without anti-PDL1, b) X-irradiation, c) X-irradiation + anti-PDL1, d) Proton irradiation, e) Proton irradiation + anti-PDL1. Error bars correspond to 1 SEM.

For MOC1 tumors, the combination with anti-PDL1 reduced the tumor growth rate compared to irradiation alone (fig.1 a-e). The reduction in growth rate η compared to irradiation alone was significantly higher for X-rays than protons (fig.3b, table 1). With addition of anti-PDL1, the growth rate reductions were significantly greater for both radiation types. The RBE_growth_ values for protons relative to X-rays show slightly higher values with anti-PDL1 compared to without anti-PDL1 (table 2). The synergistic increase in growth rate reduction was larger when combining anti-PDL1 with protons compared to X-rays (table 2).

**Table 2:**
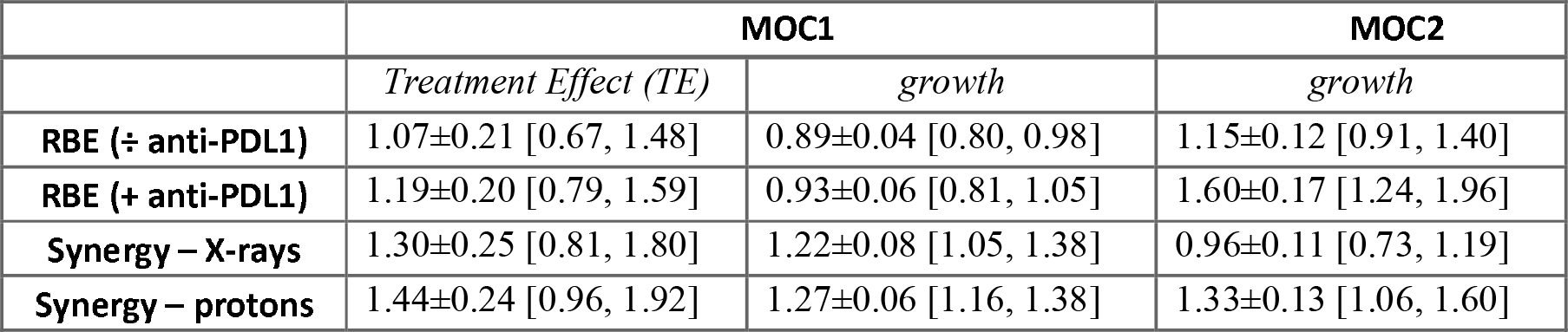
Relative biological effectiveness in terms of RBE_growth_ (MOC1 and MOC2) and RBE_TE_ (MOC1) for protons with (+) or without (÷) anti-PDL1 and synergy_growth_ (MOC1 and MOC2) and synergy_TE_ (MOC1) and when combining X- or proton irradiation with anti-PDL1 (±S.E., [95 % CI]). RBE_growth_ and synergy_growth_ calculated using data from Table 1 and eq. 2 (RBE) and eq. 3 (synergy), and RBE_TE_ and synergy_TE_ calculated using data from Figure 4 and eq. 4 (RBE) and eq. 5 (synergy).

|  | **MOC1** |  | **MOC2** |
| --- | --- | --- | --- |
|  | Treatment Effect (TE) | growth | growth |
| RBE (÷ anti-PDL1) | 1.07±0.21 [0.67, 1.48] | 0.89±0.04 [0.80, 0.98] | 1.15±0.12 [0.91, 1.40] |
| RBE (+ anti-PDL1) | 1.19±0.20 [0.79, 1.59] | 0.93±0.06 [0.81, 1.05] | 1.60±0.17 [1.24, 1.96] |
| Synergy – X-rays | 1.30±0.25 [0.81, 1.80] | 1.22±0.08 [1.05, 1.38] | 0.96±0.11 [0.73, 1.19] |
| Synergy – protons | 1.44±0.24 [0.96, 1.92] | 1.27±0.06 [1.16, 1.38] | 1.33±0.13 [1.06, 1.60] |

For MOC2 tumors, proton irradiation induced a greater growth rate reduction than X-rays, as opposed to MOC1 tumors (fig.2). Interestingly, anti-PDL1 treatment in combination with X-rays did not show any significant synergy or effect at all (fig.2a, table 2). Indeed, the growth rate reduction from X-rays alone was the same as for X-rays with anti-PDL1 (fig. 3d). However, a significantly increased growth rate reduction was found for proton irradiation together with anti-PDL1 treatment compared to proton or X-irradiation alone (fig.2b-e, table2). The RBE_growth_ values for protons relative to X-rays show higher values for MOC2 than MOC1 both with and without anti-PDL1. For MOC2 there was no synergistic increase in growth rate reduction combining X-rays with anti-PDL1, but there was a large effect with protons (table 2).

**Figure 3:**
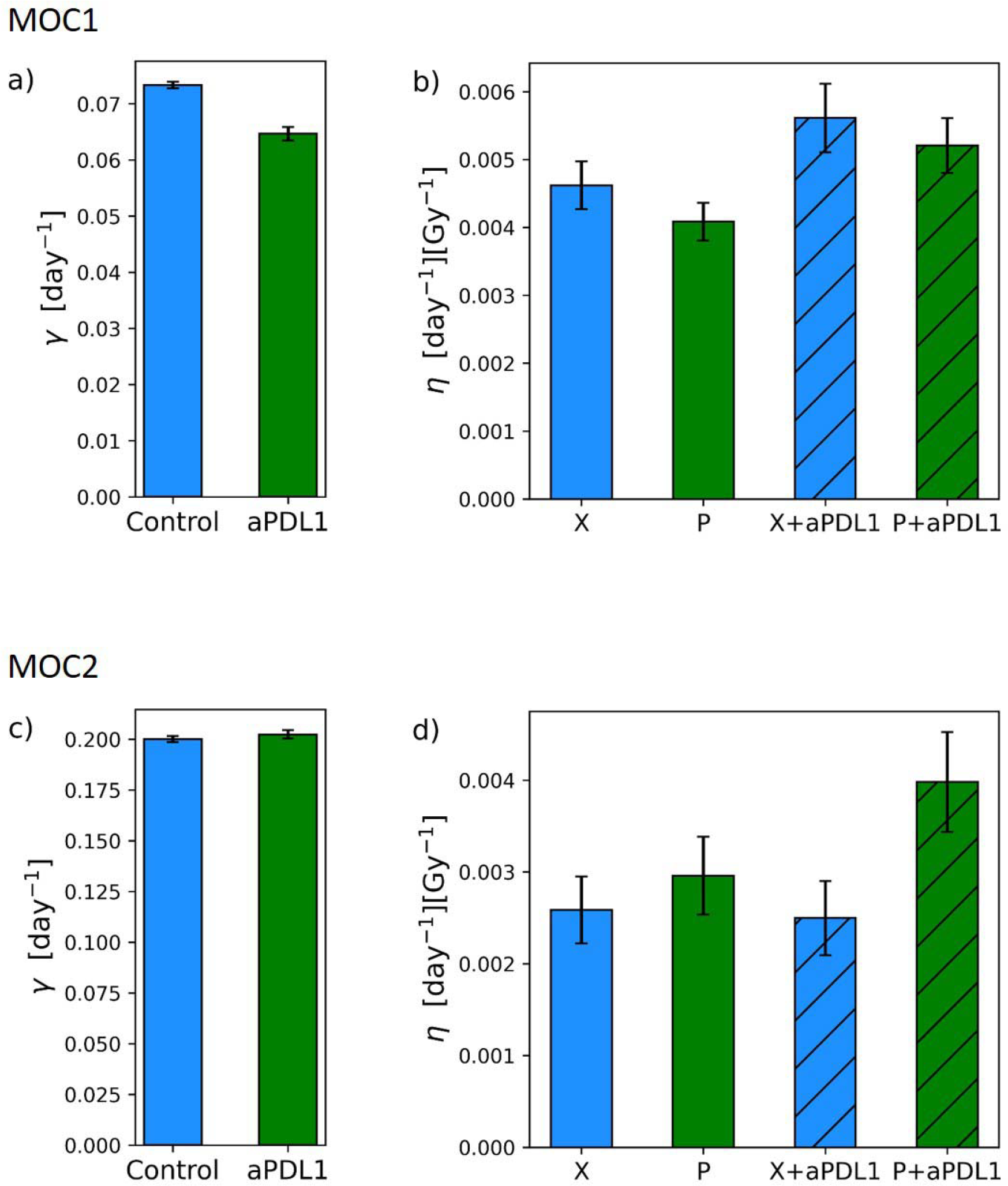
Resulting growth rate γ and growth rate reduction per dose η from model fits (eq. 1) of relative tumor volumes for MOC1 (Fig.1) and MOC2 (Fig.2). a) MOC1 growth rate γ, b) MOC1 growth rate reduction per dose η, c) MOC2 growth rate γ, b) MOC2 growth rate reduction per dose η.

MOC1 mice with complete remission (CR) on day 45 were mainly found in treatment groups receiving anti-PDL1 with doses larger than 10 Gy. For protons, CR was achieved after 10 (2/13; 15%), 15 (1/9; 11%) and 20 Gy (2/8; 25%) with anti-PDL1 treatment. For X-rays, CR was achieved for 15 (2/8; 25%) and 20 Gy (1/8; 13%) with anti-PDL1 and for 20 Gy alone (1/8; 13%). None of the MOC1-bearing mice with complete or partial remission developed metastases. The treatment effect in terms of a given animal with permanent or temporary remission (CR, PR and TR in Table 2, supplementary material) was analyzed for MOC1 tumors using logistic regression (eq. 4, Fig.4). No significant difference was observed in TD_50_ values between proton and photon radiation alone or when combined with anti-PDL1 treatment. RBE_TE_ was 1.07±0.21 without anti-PDL1 and 1.19±0.20 with anti-PDL1. The synergy_TE_ was 1.30±0.25 for X-rays and 1.44±0.24 for protons. Although the uncertainties were rather high, the RBE and synergy from the binary treatment effect outcome analysis showed the same trend as the values in table 2 estimated using the growth analysis (r=0.93).

## Discussion

There is great interest in combining radiotherapy and PDL1 blockade, as the efficacy of either treatment is limited for many diseases. Still, it is not clear how to optimize the combination treatment, as the outcome depends on radiation type, dose, and fractionation, among others [25]. To our knowledge, we present the first study assessing the efficacy of combining proton therapy with an immune checkpoint inhibitor. We have shown that the MOC1 and MOC2 tumors responded very differently to protons and X-rays with or without anti-PDL1-treatment. MOC1 tumors grew slower, were more radiosensitive, and showed less metastases than MOC2 tumors. For MOC1, the short-term tumor response appeared to be higher for X-ray monotherapy compared to proton monotherapy. For immunotherapy, a 20 % reduction in MOC1 growth rate was found without irradiation. When combined with radiotherapy, a possible synergistic effect was seen for both protons and X-rays. For MOC2, a slightly higher short-term tumor response was observed for protons compared to X-rays. Immunotherapy alone had no effect on MOC2 tumor growth, but in combination with radiotherapy, protons gave a significant synergistic effect while this was not the case for X-rays.

The higher efficacy of X-rays compared to protons for MOC1 tumors at day 19 can be quantified from the growth analysis to be RBE_growth_ = 0.83±0.03 without anti-PDL1. A proton RBE of ∼0.8 has been reported previously for tumor growth delay of NFSa fibrosarcoma cells transplanted in C3H/He mice [26]. However, using the treatment effect on day 45 (fig. 4) as end point gave RBE_TE_ = 1.07±0.21, which is closer to the commonly used RBE of 1.1 although with large uncertainties. For MOC2 tumors, protons had a slightly higher effect than X-rays (not significant) when given alone. Combining irradiation with anti-PDL1 treatment increased RBE_growth_ for both MOC1 and MOC2, where the impact was greatest for the latter tumor model. Both without and with anti-PDL1 the RBE_growth_ for MOC2 tumors was significantly higher than for MOC1, indicating that RBE is affected by adjuvant use of immunotherapy and by tumor characteristics. From H&E-stained tumor sections (supplementary file), we found that MOC2 tumors were largely undifferentiated while MOC1 was a moderately to highly differentiated oral squamous cell carcinoma. Thus, the most differentiated tumor (MOC1) had the lowest RBE, which is in line with the findings by Glowa et al. who found that RBE for carbon ion therapy decreased with tumor differentiation for rat prostate cancers [27]. As in the present study, the increase in RBE predominantly resulted from a decrease in photon effect with increasing differentiation while the variation in treatment response between the rat cell lines to carbon ions was smaller.

**Figure 4:**
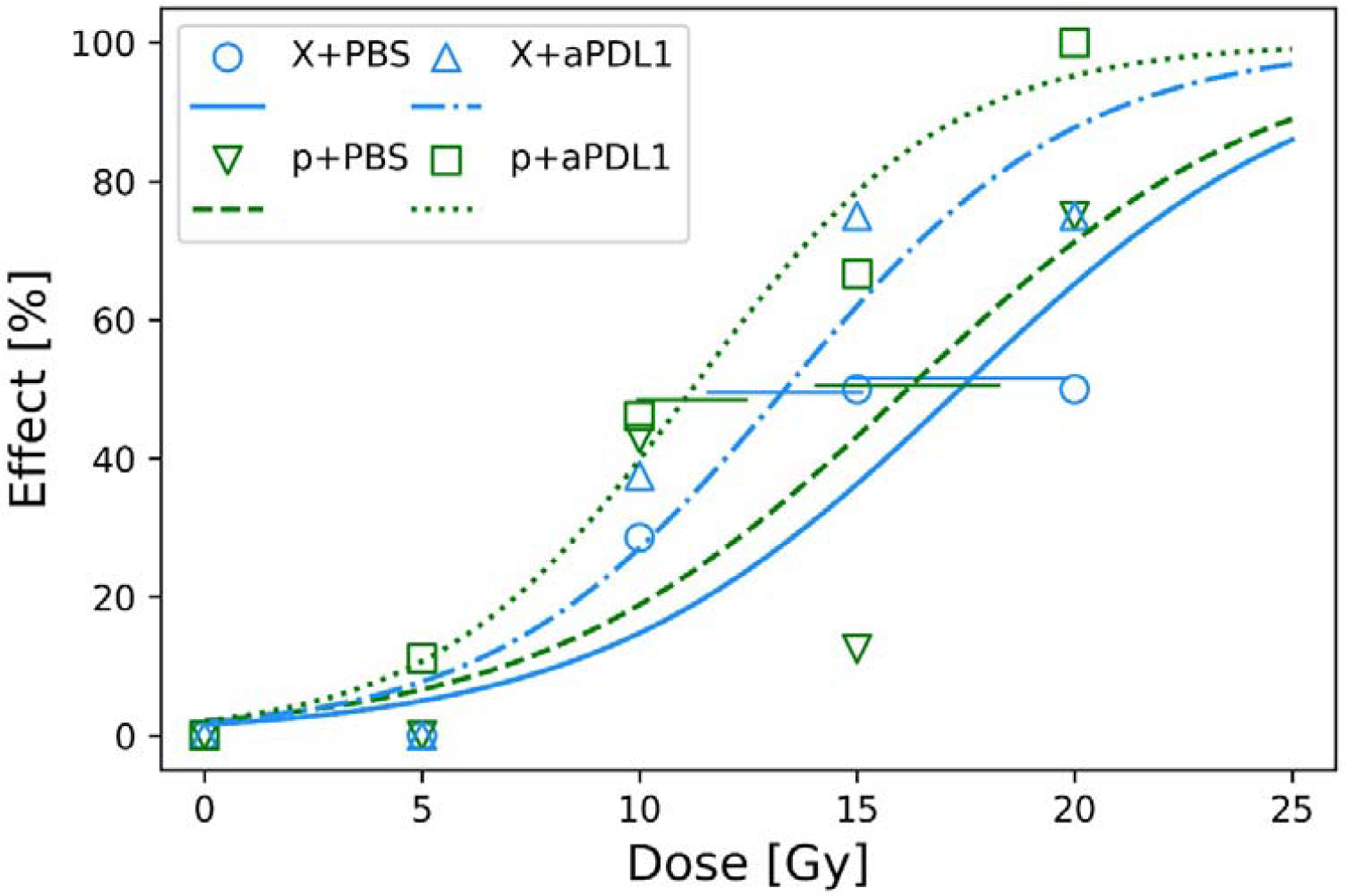
Tumor effect in terms of fraction of animals not showing progressive disease after treatment for MOC1 tumors evaluated at day 45. Points show data, while lines correspond to logistic regressions on data for respective treatment groups with estimated SEM in TD_50_ as horizontal bars.

The rationale for combining irradiation with immune checkpoint inhibitors is that inhibition of immune checkpoints only has an effect if the immune system recognizes the tumor cells as foreign. Radiation has been shown to induce immunogenic cell death, which activates anti-tumor cytotoxic T cells. The two MOC cell lines were chosen due to their different immunogenicity with MOC1 being very immunogenic, and therefore expected to respond to immune checkpoint inhibition without irradiation. MOC2, on the other hand, is less immunogenic and in addition very aggressive [22]. Consistent with these characteristics, we found a significant reduction in MOC1 tumor growth rate in mice treated with anti-PDL1 alone. The growth rate of untreated MOC2 tumors was much higher than for MOC1 and was not affected by anti-PDL1 treatment. In the study by Judd et al. comparing growth of MOC1 and MOC2 tumors in immunodeficient RAG2–/– mice and WT C57Bl/6 mice, MOC2 tumors grew at the same rate in immunodeficient and WT mice and generally faster than MOC1 tumors. Conversely, MOC1 tumors grew slower in WT than in immunodeficient mice [21]. In addition, an elevated CD8+ T cell infiltration in MOC1, but not MOC2 tumors, was observed, indicating that the MOC1 tumor growth was slowed down by the immune system [21].

A significantly lower tumor growth rate of MOC2 tumors was found for protons in combination with anti-PDL1 compared to proton therapy alone, with a synergy_growth_ of 1.33±0.13. Interestingly, combining anti-PDL1 with X-rays did not change the growth rate compared to X-rays alone for MOC2 tumors. Since the MOC2 tumors were not immunogenic without irradiation, the synergistic effect with protons and anti-PDL1, but not X-rays, suggests that protons are more efficient in inducing an immunogenic response. For the already immunogenic MOC1 tumors, we observed a synergistic effect for both X-rays and protons combined with anti-PDL1. This could either indicate that the irradiation improved the already present immunogenic response or that it decreased immunosuppressive mechanisms. Since only protons showed synergy with anti-PDL1 for the less immunogenic MOC2 tumors and protons also showed a higher synergy than X-rays for MOC1, one could speculate whether protons are superior in inducing immunogenic responses, while X-rays may increase tumor delay through different mechanisms. This is partly substantiated by a study comparing radiation-induced immunogenicity for X-rays and carbon ions [28]. In that study, carbon ions gave both high levels DAMPs in the form of HMGB1 and low levels of immunosuppressive cytokines while X-rays only gave increased HMGB1 levels [28].

For MOC1 tumors, only a few of the treated animals developed metastases, and these mice had only temporary tumor remission. Thus, for the MOC1 cell line, metastases seemed to be a consequence of high tumor burden. All animals with MOC2 tumors developed metastases and even the mice with complete tumor response had to be euthanized preventing long term follow-up. This implies that metastases also developed during anti-PDL1 treatment and thus only a local, but not a systemic response was seen for MOC2 tumors. This would suggest that MOC2 tumors develop metastases with immunosuppressive characteristics, which do not respond to antitumor immune signals, even in the animals where the tumor shows a synergistic response to radiation and anti-PDL1. It also indicates that inducing an immunogenic response and blocking immune checkpoints are not sufficient to induce a response in MOC2 metastases. Thus, MOC2 appears to be an excellent model for future studies aiming at developing treatment strategies for aggressive H&N tumors that are resistant to radiation- and immune therapy.

In conclusion, we found synergistic effects from combining protons and immunotherapy in the less immunogenic MOC2 tumor model with poorly differentiated squamous cell carcinoma cells, which was not evident for X-rays. It was not possible to detect a systemic effect, as all animals bearing MOC2 tumor developed metastases. In the immunogenic MOC1 tumor model with well-differentiated tumor tissue, anti-PDL1 had an effect alone, confirming that the tumor was already immunogenic. In this tumor, a synergistic effect of anti-PDL1 and irradiation was seen for both protons and X-rays, and high tumor effect prevented the development of metastases. Thus, protons appear to be a better choice when combined with ICI, but our findings warrant further validation and mechanistic studies.

