## Supplementary Material for "Synergistic effects of combing proton- or X-irradiation with anti-PDL1 immunotherapy in two murine oral cancers"

*Table 1: Number of mice in each treatment group.*

| **Treatment** | | | | |
|---|---|---|---|---|
| **Modality** | **IP Injections** | **Dose (Gy)** | **MOC1** | **MOC2** |
| | PBS | 0 | 31 | 38 |
| | aPDL1 | 0 | 8 | 17 |
| **X-rays** | PBS | 5 | 7 | - |
| | | 10 | 7 | 14 |
| | | 15 | 8 | - |
| | | 20 | 8 | 13 |
| | | 30 | - | 15 |
| | aPDL1 | 5 | 9 | - |
| | | 10 | 8 | 14 |
| | | 15 | 8 | - |
| | | 20 | 8 | 14 |
| | | 30 | - | 15 |
| **Protons** | PBS | 5 | 8 | - |
| | | 10 | 14 | 13 |
| | | 15 | 8 | - |
| | | 20 | 8 | 12 |
| | | 30 | - | 14 |
| | aPDL1 | 5 | 9 | - |
| | | 10 | 13 | 14 |
| | | 15 | 9 | - |
| | | 20 | 8 | 12 |
| | | 30 | - | 14 |

*Table 2: Treatment effect on day 45 for MOC1 tumors. All mice were categorized as either Progressive Disease (PD), Temporary Remission (TR), Partial Remission (PR) or Complete Remission (CR) as defined in the methods section.*

| **Treatment** | | **Treatment Effect (TE)** | | | | |
|---|---|---|---|---|---|---|
| **Modality** | **IP Injections** | **Dose (Gy)** | **PD** | **TR** | **PR** | **CR** |
| | PBS | 0 | 31/31 | | | |
| | aPDL1 | 0 | 8/8 | | | |
| **X-rays** | PBS | 5 | 7/7 | | | |
| | | 10 | 5/7 | 2/7 | | |
| | | 15 | 4/8 | 3/8 | 1/8 | |
| | | 20 | 4/8 | 3/8 | | 1/8 |
| | aPDL1 | 5 | 9/9 | | | |
| | | 10 | 5/8 | 2/8 | 1/8 | |
| | | 15 | 2/8 | 3/8 | 1/8 | 2/8 |
| | | 20 | 2/8 | 1/8 | 4/8 | 1/8 |
| **Protons** | PBS | 5 | 8/8 | | | |
| | | 10 | 8/14 | 5/14 | 1/14 | |
| | | 15 | 7/8 | 1/8 | | |
| | | 20 | 2/8 | 5/8 | 1/8 | |
| | aPDL1 | 5 | 8/9 | 1/9 | | |
| | | 10 | 7/13 | 4/13 | | |
| | | 15 | 3/9 | 4/9 | 1/9 | 1/9 |
| | | 20 | | 5/8 | 1/8 | 2/8 |

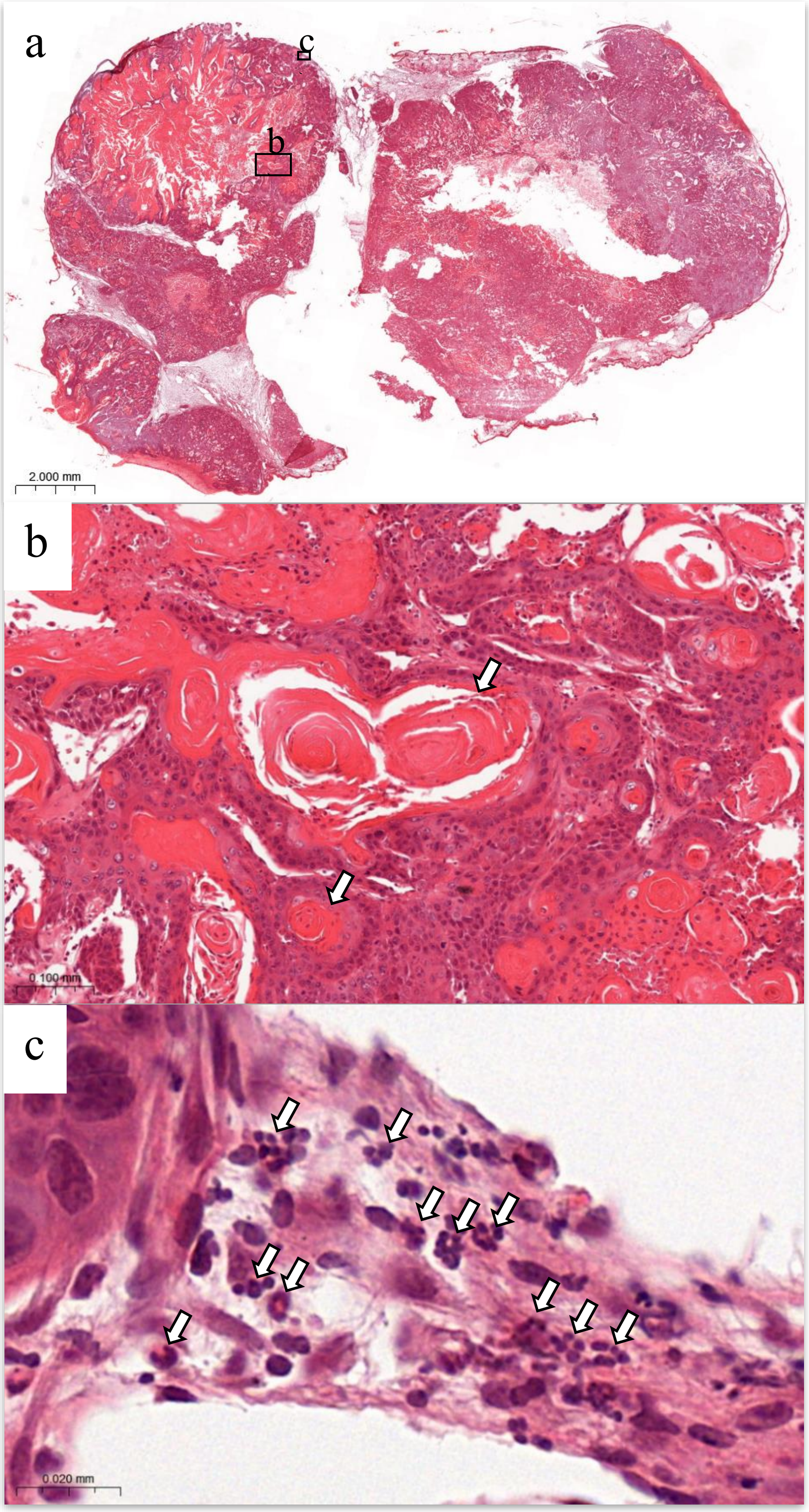


*Figure 1: Histological picture of hematoxylin and eosin stained (as described in the method below) untreated MOC1 tumor tissue on day 41. a) Section of whole tumor is shown. The tumor is cell rich with focal areas of keratinization (b). Small bands/areas of connective tissue are seen, both within and in the periphery of the tumor tissue. Here, a few chronic inflammatory cells are identified. Neutrophil granulocytes are present, particularly in the peripheral part of the tumor (c). b) Example of area with keratinization. c) Examples of neutrophil granulocytes in the peripheral part of the tumor, marked with arrows.*

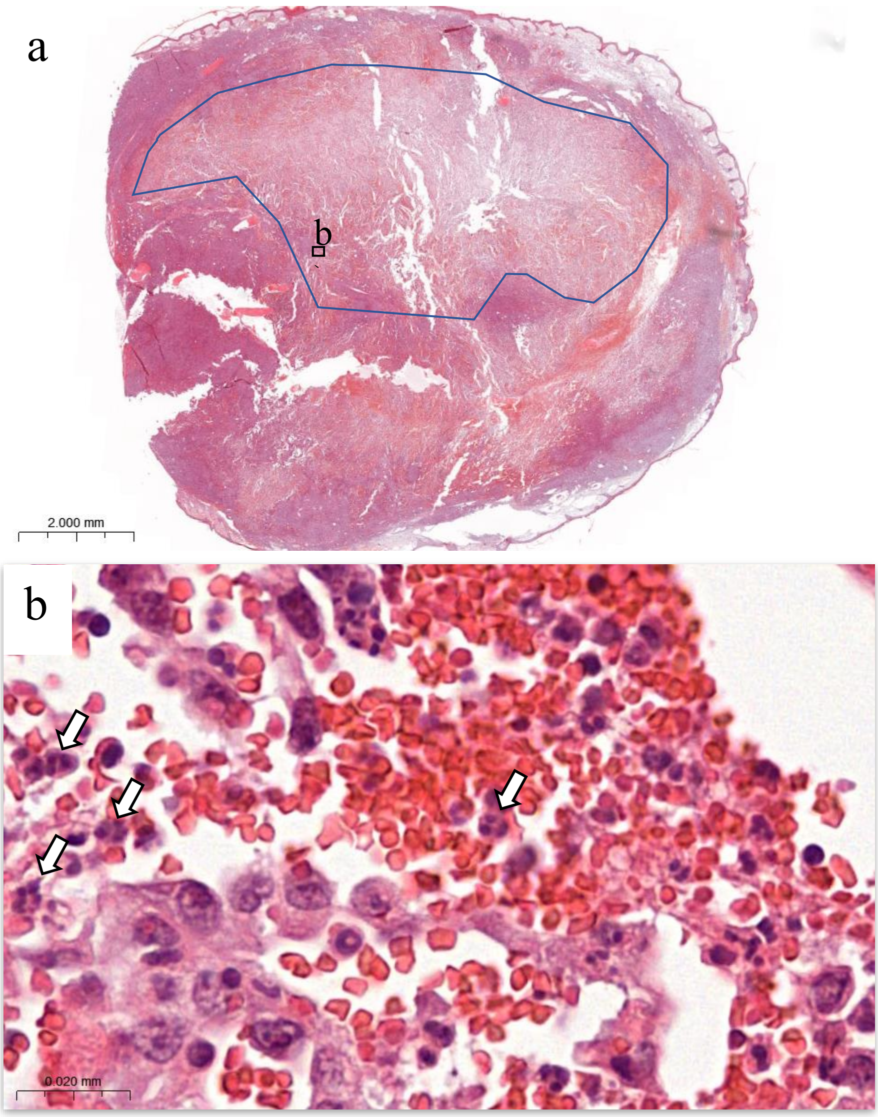


*Figure 2: Histological picture of a hematoxylin and eosin stained (as described in the method below), poorly differentiated untreated MOC2 tumor on day 12 after treatment start. a) Section of whole tumor is shown. The tumor tissue is disintegrated and areas of partly necrotic and necrotic tumor tissue (example marked with blue line) with infiltration of neutrophils dominates (b). Extravascular erythrocytes are seen throughout the tumor tissue. b) Necrotic tissue with examples of neutrophils marked with arrows. Extravascular erythrocytes can be seen.*

**Materials and methods - Hematoxylin and eosin staining**

Four micron thick paraffin sections were mounted on object slides, dried for 1 h at 60 °C, deparaffinized in two changes of xylene for 5 min each and rehydrated in two changes of absolute ethanol and 96 % ethanol, and finally 70 % ethanol for 2 min each before washing in tap water for 5 min. Nuclei were stained for 1 min with hematoxylin (Shandon, Thermo Fisher Scientific, Waltham, MA, USA), washed in tap water for 5 min and blued in 0.25 % w/v hexamethylenetetramine (Prolabo, Fontenay-sous-Bois , F) for 3 min. After washing in tap water for 5 min, cytoplasm was stained using 0.5 % w/v Eosin-Yellow (Chroma, Münster, D) for 4 min before the slides were rinsed in tap water, quickly dehydrated through graded alcohols, cleared in xylene and cover glass mounted with Histokitt (Karl Hecht, Sondheim, D).
